# Group-Level Analysis of Induced Electric Field in Deep Brain Regions by Different TMS Coils

**DOI:** 10.1101/786459

**Authors:** Jose Gomez-Tames, Atsushi Hamasaka, Akimasa Hirata, Ilkka Laakso, Mai Lu, Shoogo Ueno

**Author notes:** Corresponding Author: Jose Gomez-Tames.

## Abstract

Deep transcranial magnetic stimulation (dTMS) is a non-invasive technique used in treating depression. In this study, we computationally evaluate group-level dosage during dTMS with the aim of characterizing targeted deep brain regions to overcome the limitation of using individualized head models to characterize coil performance in a population.

We use an inter-subject registration method adapted to deep brain regions that enable projection of computed electric fields (EFs) from individual realistic head models (*n* = 18) to the average space of deep brain regions. The computational results showed consistent group-level hotspots of the EF in deep brain region with intensities between 20%-50% of the maximum EF in the cortex. Large co-activation in other brain regions was confirmed while half-value penetration depth from the cortical surface was smaller than 2 cm. The halo figure-8 assembly and halo circular assembly coils induced the highest EFs for caudate, putamen, and hippocampus.

Generalized induced EF maps of deep regions show target regions despite inter-individual difference. This is the first study that visualizes generalized target regions during dTMS and provides a method for making informed decisions during dTMS interventions in clinical practice.

## 1. INTRODUCTION

Transcranial magnetic stimulation (TMS) (Barker et al. 1985) is a non-invasive technique for brain neurostimulation that is used in clinical diagnosis and to understand brain functions in neuroscience. Recently, repetitive TMS has been used in treating depression (Fitzgerald et al. 2006). During TMS, a high-intensity pulsed current is applied to a coil placed on the head, and its generated magnetic field induces eddy currents in the brain through the Faraday law. The induced current/electric field (EF) can activate specific cortical targets.

Recent studies (Levkovitz et al. 2009) indicate that deep brain regions are involved in psychoneurotic diseases such as depression. Similarly, structures related to the reward systems include the nucleus accumbens (NAcc), ventral tegmental area, and amygdala (Nemeroff 2002; Russo and Nestler 2013). These structures are located 4.5–6.5 cm from the scalp, and their stimulation can lead to antidepressant effects in deep brain stimulation (DBS) (Berton and Nestler 2006; Schlaepfer et al. 2008). Deep TMS (dTMS) has been proposed as an alternative non-invasive procedure (Levkovitz et al. 2009; Tendler et al. 2016; Kedzior et al. 2018) and has been approved by the US Food and Drug Administration (FDA) for treatment of obsessive compulsive disorder. To stimulate the target areas located in deep regions beneath the scalp, different coils have been proposed for realizing stimulation more efficiently than with the conventional figure-8 coil (Ueno et al. 1988). These include the H-coil (Roth et al. 2002), halo figure-8 assembly (HFA) coil (Lu and Ueno 2015), halo circular assembly (HCA) coil (Crowther et al. 2011), double cone coil (Lontis et al. 2006), and H7 coil (Popa et al. 2019). The purpose of designing these new coils is to overcome the trade-off between depth and dispersion (Deng et al. 2013; Lu and Ueno 2017), in which reaching deeper brain structures implies a wider EF spread on the cortical and subcortical regions in the form of co-activation (Guadagnin et al. 2016). Previous dTMS computational study evaluated the EF strength in a sphere that mimicked the head model anatomical shape (Deng et al. 2014). The findings from these studies are useful for understanding the fundamental difference of the EF in deep regions. Subsequent studies evaluated the individual effects of EF in the different subcortical regions using detailed head model subjects (Fiocchi et al. 2016; Guadagnin et al. 2016; Lu and Ueno 2017; Parazzini et al. 2017; Samoudi et al. 2018). However, EF effects derived from individualized models cannot always be generalized to a group of subjects as the EF in the brain is distorted by the brain’s complicated structure, resulting in large variability (Gomez-Tames et al. 2018). To overcome this limitation, inter-subject registration methods have been applied for cortical regions to investigate group-level effects of TMS parameters and coil design in previous studies (Iwahashi et al. 2017; Kutsuna et al. 2017; Mikkonen et al. 2018). One element that has being missing is the group level analysis in deep brain regions that is required for coil performance evaluation in dTMS studies.

The purpose of this study is to evaluate the induced EF in each target area of deep brain regions at a group level to characterize coil performance. To achieve it, we extend proposed an inter-subject registration method on deep brain regions. This approach also permitted the evaluation of trade-off between depth and dispersion, co-activation, and optimization of coil location.

## 2. METHOD AND MODEL

### A. Anatomical models of the human brain

Eighteen anatomical head models with a resolution of 0.5 mm were obtained from our previous study (Laakso et al. 2015). The models were constructed from T1- and T2-weighted images acquired from a magnetic resonance image scanner (available on: http://hdl.handle.net/1926/1687). FreeSurfer image analysis software (Dale et al. 1999; Fischl 2012) was used to reconstruct the surfaces of the gray, white matter, cerebellum gray matter, cerebellum white matter, and deep brain regions (brainstem, thalamus, caudate, putamen, pallidum, hippocampus, amygdala, NAcc). The tissue other tissues compartments were segmented by semiautomatic methods (Laakso et al. 2015, 2016) into the following: skin, fat, muscle, blood, outer skull, inner skull, intervertebral disc, ventricular cerebrospinal fluid, ventral diencephalon, meninges, mucous membrane, and dura, as shown in Fig. 1(a). Note that the cerebrospinal fluid was the volume inside the skull that was not classified as nervous tissue or blood explicitly.

**Figure 1.**
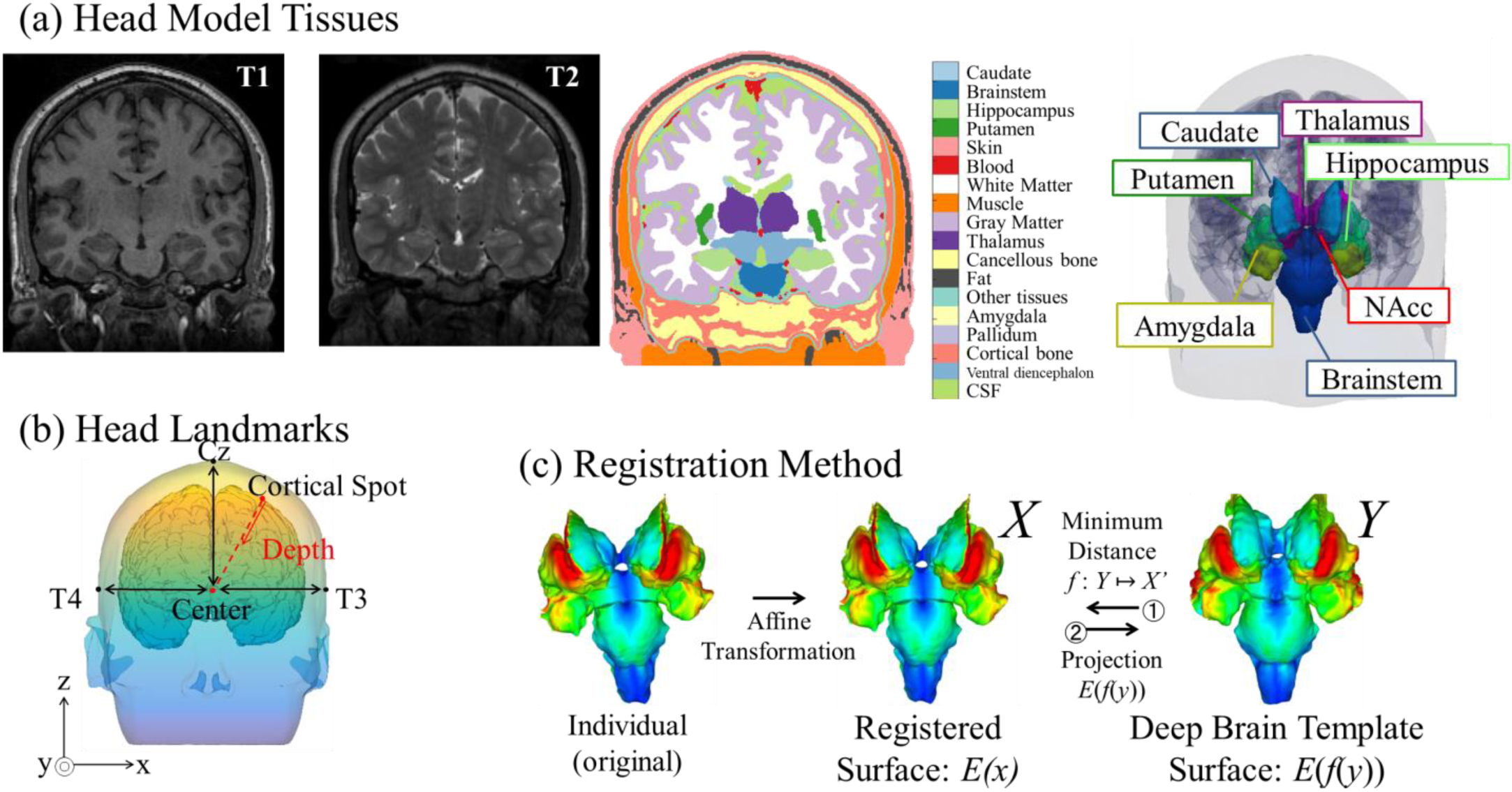
(a) Head model segmentation including the seven deep brain regions under consideration. (b) Definition of the depth and focality metrics at the “center” of the brain. (c) Registration method of the EFs on the deep-brain-region surfaces to the standard deep brain template.

### B. Computational methods

A volume conductor model was used to compute induced EFs in the head models. The magneto-quasi-static approximation applies to the 10-kHz frequency band, and we assume that the displacement current is negligible when compared with the conduction current (Hirata et al. 2013). In addition, the induced current does not perturb the external magnetic field. The induced scalar potential *φ* is given by:

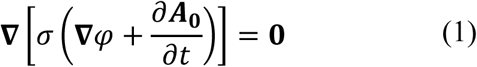

where ***A***_0_ and *σ* denote the magnetic vector potential of the applied magnetic field and tissue conductivity, respectively. The induced EF was calculated from the gradient of the scalar potential by the following expression:

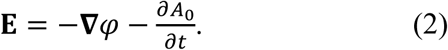

The Eq. (1) was solved numerically using the finite-element method (FEM) with first-order 0.5 mm × 0.5 mm × 0.5 mm cubical elements (Laakso and Hirata 2012). The electric conductivity of head tissues was assumed to be linear and isotropic. As shown in Table 1, a different tissue conductivity was assigned to each tissue based on the fourth-order Cole-Cole model at 10 kHz (Gabriel et al. 1996). The coil current was fixed to 1 A for all simulations. Associated numerical errors were marginal considering the model resolution (Gomez-Tames et al. 2017, 2019b) and experimental verifications (Laakso et al. 2017; Aonuma et al. 2018; Mikkonen et al. 2018). In addition, averaging over multiple subjects further reduces the error in the mean EF strength as follows in the next subsection.

**TABLE 1.** Electric conductivity values of the head model tissues

| Tissue/ bodily fluid | Conductivity<br>[S/m] |
| --- | --- |
| Amygdala | 0.12 |
| Blood | 0.7 |
| Bone (cancellous) | 0.04 |
| Bone (cortical) | 0.01 |
| Brainstem | 0.07 |
| Caudate | 0.12 |
| Cerebellum gray matter | 0.1 |
| Cerebellum white matter | 0.07 |
| Cerebrospinal fluid | 1.8 |
| Dura | 0.2 |
| Fat | 0.08 |
| Gray matter | 0.12 |
| Hippocampus | 0.12 |
| Intervertebral disc | 0.1 |
| Meninges | 0.2 |
| Mucous membrane | 0.1 |
| Muscle | 0.2 |
| Nucleus accumbens | 0.12 |
| Pallidum | 0.12 |
| Putamen | 0.12 |
| Skin | 0.08 |
| Thalamus | 0.12 |
| Ventral diencephalon | 0.07 |
| Ventricular cerebrospinal<br>fluid | 1.8 |
| Vitreous humor | 1.6 |
| White matter | 0.07 |

### C. Registration method

Surface data in the seven deep brain structures (brainstem, caudate, putamen, hippocampus, amygdala, NAcc, and thalamus) were registered to the deep regions of the MNI ICBM 2009a standard template (Fonov et al. 2009, 2011). For each deep brain region, we took the surface mesh and the same surface for the MNI template and used an iterative closest point method (part of the Visualization Toolkit (VTK)) to get an affine registration between the surfaces. For each point *y* in the template surface of a deep region *Y*, we found the closest point *x* in the registered individual deep region *X* by the means of the minimum Euclidian distance (*f*: *Y* → *X*). If *E* was the EF magnitude from an individual deep region, the template EF at y was calculated by *E*(*f*(*y*)) (Gomez-Tames et al. 2019a). The process was repeated for each deep region and summarized in Fig. 1(c). In addition to deep brain regions, the surface EFs were registered to the surface of the brain of the MNI ICBM 2009a standard template using a previously described registration procedure for co-activation analysis in Fig. 5(b) (Laakso et al. 2015).

### D. Group-level EF characterization

To compare the characteristics of dTMS coils, the group level EF was the average of the registered EF strengths (absolute value) in the standard brain space between all the subjects for the seven deep brain regions to minimize inter-individual effects. The inter-individual effect was quantified by the relative standard deviation of the EFs. In addition, the EFs were normalized with the maximum value in the cerebral cortex to facilitate comparison between coils despite differences in the coil design and number of turns. To mitigate numerical artifacts derived from computing the EF with the voxel model at the surface of the CSF–brain boundaries (Reilly and Hirata 2016), we used the 99.9th percentile value of the EF as the maximum value (Gomez-Tames et al. 2017).

It is possible that some points on the individual surface were not assigned to the standard deep brain template with the potential loss of hotspots or EF information due to the minimum Euclidian distance criteria. We defined a metric of the registration error to investigate the potential information loss. The registration error for each subject was the difference between two EF distributions: (i) *E*(*x*) which is the original EF in the individual deep regions and (ii) *E* (*f*(*g*(*x*))) which is the registration of *E*(*f*(*y*)) back to the individual deep region using the Euclidean distance (*g*: *X* → *Y*) as follows

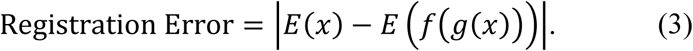

This study adopted the EF strength to describe regions with a high possibility of stimulation. The rationale is that even in the cerebral cortex where there is a highly uniform orientation of the pyramidal neurons relative to the cortical surface, a consensus of the most appropriate EF direction has not been established (Fox et al. 2004; Bungert et al. 2016; Laakso et al. 2017). In the case of the deep brain regions, the orientation of the majority of the neurons, which is important to determine the most effective EF direction, is not clear except for hippocampus.

### E. Coil positioning scenarios and modelling

Group-level EF comparison of seven coils was conducted for two scenarios. In the first one (subsection 3.A to 3.C), all the coils (except H1) were centered at the same position (cz) in all subjects to reduce intra-subject anatomical variations of the tissues underneath the coil (e.g., the distance to deep regions). The H1 coil was placed on the frontal region (around afz) and adjusted to fit the head geometry. In the second scenario (subsection 3.D), coils were compared using optimized coil locations based on the 10-10 international EEG system. Locations were omitted if the windings crossed the head model. All the coils were optimized using a medial-lateral orientation.

A schematic of the seven coils is shown in Fig. 2. Fig. 2(a) shows a circular coil whose inner and outer radii are 35 and 45 mm, respectively. The number of turns is 14. Fig. 2(b) shows a commonly used figure-8 coil whose inner and outer radii are 23.5 and 48.5 mm, respectively. The number of turns is nine. Fig. 2(c) shows the H coil based on a previous study (Lu and Ueno 2017). The coil has 13 windings and consists of a base and return parts. A base part is oriented along the anterior-posterior axis on the left hemisphere, thus stimulating neuronal pathways along this axis. The return part directs the return currents on the right hemisphere. Fig. 2(d) shows a halo figure-8 assembly coil, which combines the figure-8 coil shown in Fig. 2(b) and the halo coil. The halo coil radii are 138 and 144 mm, and their position is 100 mm below the figure-8 coil. The number of turns is five. Fig. 2(e) shows the halo circular assembly coil, which combines the circular and halo coils. Fig. 2(f) shows the double cone coil, in which the angle formed between the two circular coils is 95 degrees (instead of 180 degrees), as with the figure-8 coil. The inner and outer radii of the circular coils are 48 and 65 mm, respectively, and the number of turns is nine. Fig. 2(g) shows the H7 coil, consisting of two adjacent wings fixed at a relative angle of 90 degrees. Each wing consists of two layers of concentric elliptical windings with major axis ranging from 75–140 mm, and minor axis ranging from 70−125 mm; and the number of turns is four.

**Figure 2.**
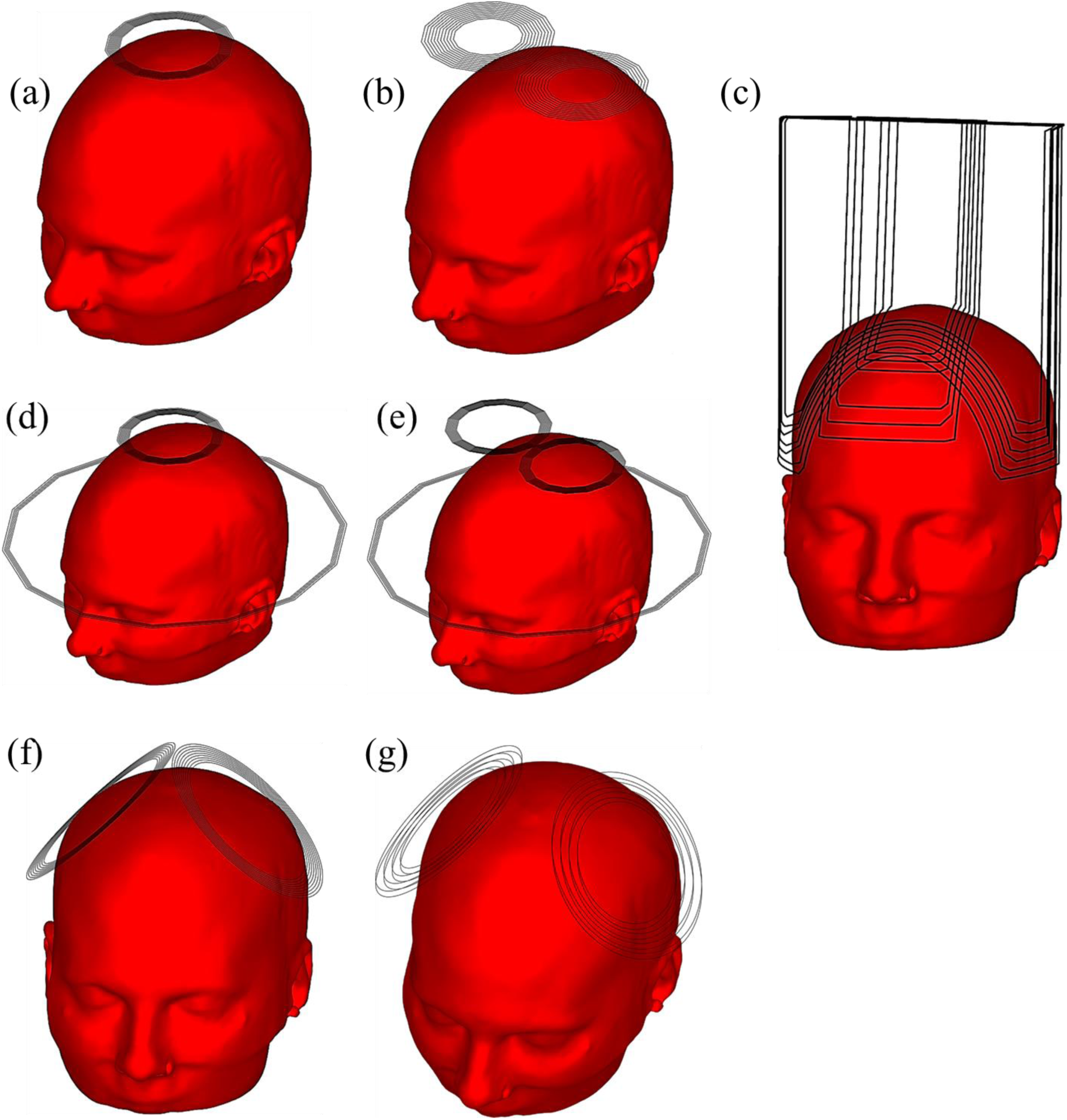
Seven coils under comparison: (a) circular, (b) figure-8, (c) H, (d) HCA, (e) HFA, (f) double cone, and (g) H7 coils for medial-lateral orientation.

### F. EF penetration and spread (co-activation)

The penetration depth and spread of the EF from the cortex were examined. Specifically, the spread to the “depth” direction was quantified by examining the penetration and volume of the EF. The “depth” direction was defined using the same method as in (Guadagnin et al. 2016). In brief, the “center” of the brain was defined under cz at the height of T3 and T4 by the 10-20 EEG system. In addition, we determined the location of the maximum EF strength on the cortex. Note that the position of the original maximum value and 99.9^th^ percentile value was nearly the same. The direction from the “cortical spot” to the “center” was defined as the “depth” direction, as shown in Fig. 1(b).

Moreover, the half-value spread metric (inverse of focality) was used to indicate the spread in the “depth” direction as follows:

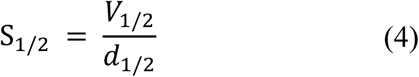

where *V*_1/2_ is the half-value volume defined as the volume in the brain region in which the EF is equal to or greater than 50% the value of EF on the “cortical spot”. The term *d*_1/2_ is the half-value depth corresponding to the maximum depth in which the EF is equal to or greater than 50% the EF value on the “cortical spot” We calculated the mean value and standard deviation of *d*_1/2_, *V*_1/2_, and S_1/2_ for 18 models.

## 3. COMPUTATIONAL RESULTS

### A. EF registration

Fig. 3(a) illustrates the effect of individual variability on the EF distribution of the seven deep brain regions for the case of HCA coil. For instance, we can observe a distinct EF pattern in the amygdala for subject 10 and subject 11. To investigate group-level EF effects, the individual deep EFs is transformed to the standard deep brain template EFs, as shown in Fig. 3(b). Visual inspection shows an adequate matching between individual and template EFs distributions. Also, fig. 3(c) confirms the good matching with a median registration error of 2.6 mV/m (individual deep regions range: 1.3-4.2 mV/m) or normalized median error of 0.7%.

**Figure 3.**
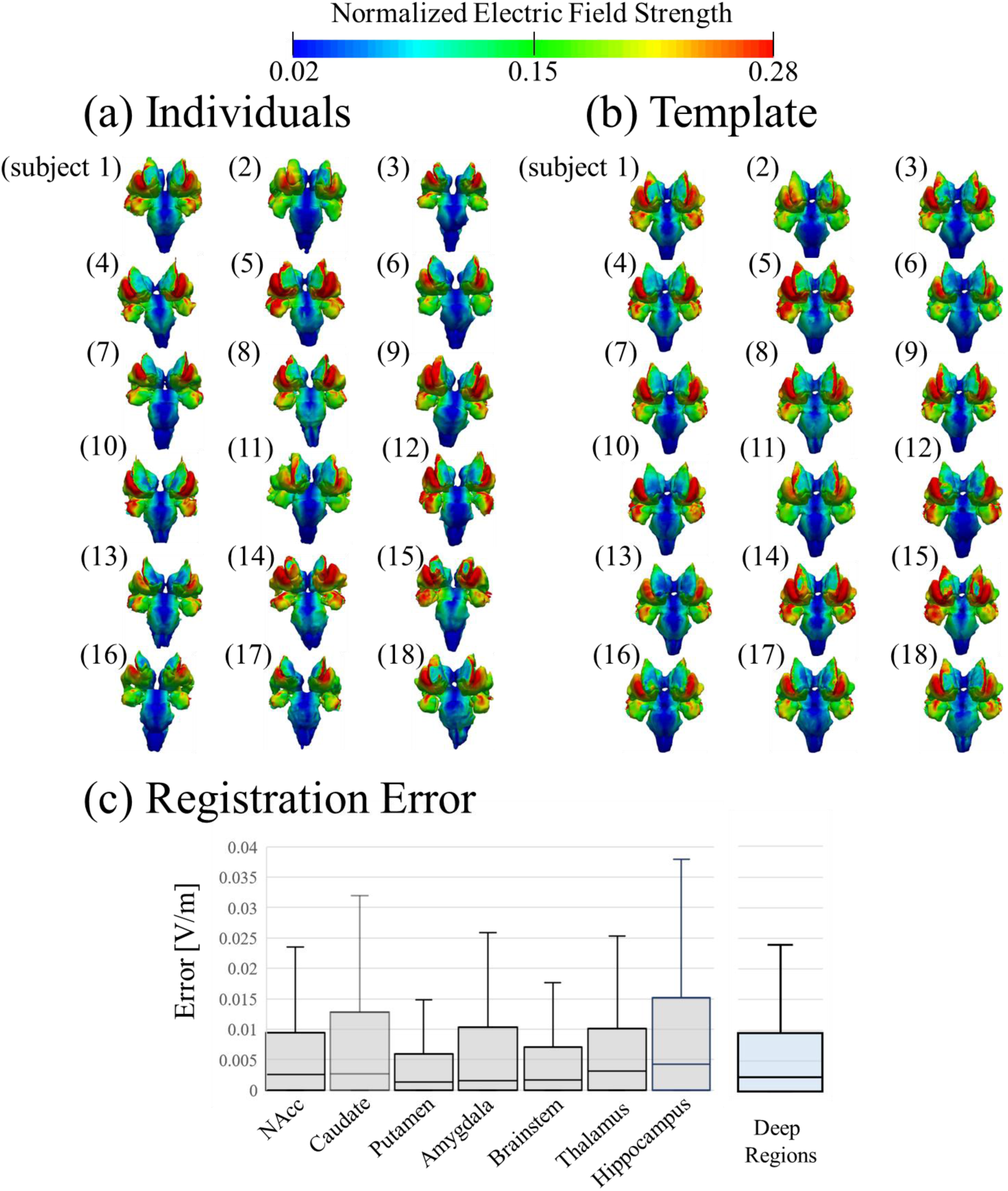
Individualized EF in the deep brain regions for the HCA coil centered in cz position (values are normalized by the maximum cortical EF.). (a) EF distribution of each head model. (b) EF distribution projected on the standard space. (c) Registration error for each deep brain region and whole deep regions for HCA coil (*n* = 18).

### B. Group-level EF distribution difference between coils

Fig. 4(a) shows the group-averaged (*n* = 18) EF distribution on the standard space of the seven deep brain regions for all coils centered at the same position (except H1 in the frontal region) to exclude intra-subject differences (i.e., distance to deep regions). The HFA and HCA induced the highest EFs in the deep regions in relationship to cortical EFs. In contrast, circular and figure-8 coil produced weaker EFs. High EFs were distributed mostly in the caudate and putamen for all coils. The third structure with higher EFs was variable among the coils (e.g., hippocampus for HFA and HCA and brainstem for figure-8 coil). In addition, Fig. 4(b) shows the standard deviation of the induced EF for each coil. The caudate region presents the larger variation. The group-level hotspots of the EF in deep brain regions were 20-40%.

**Figure 4.**
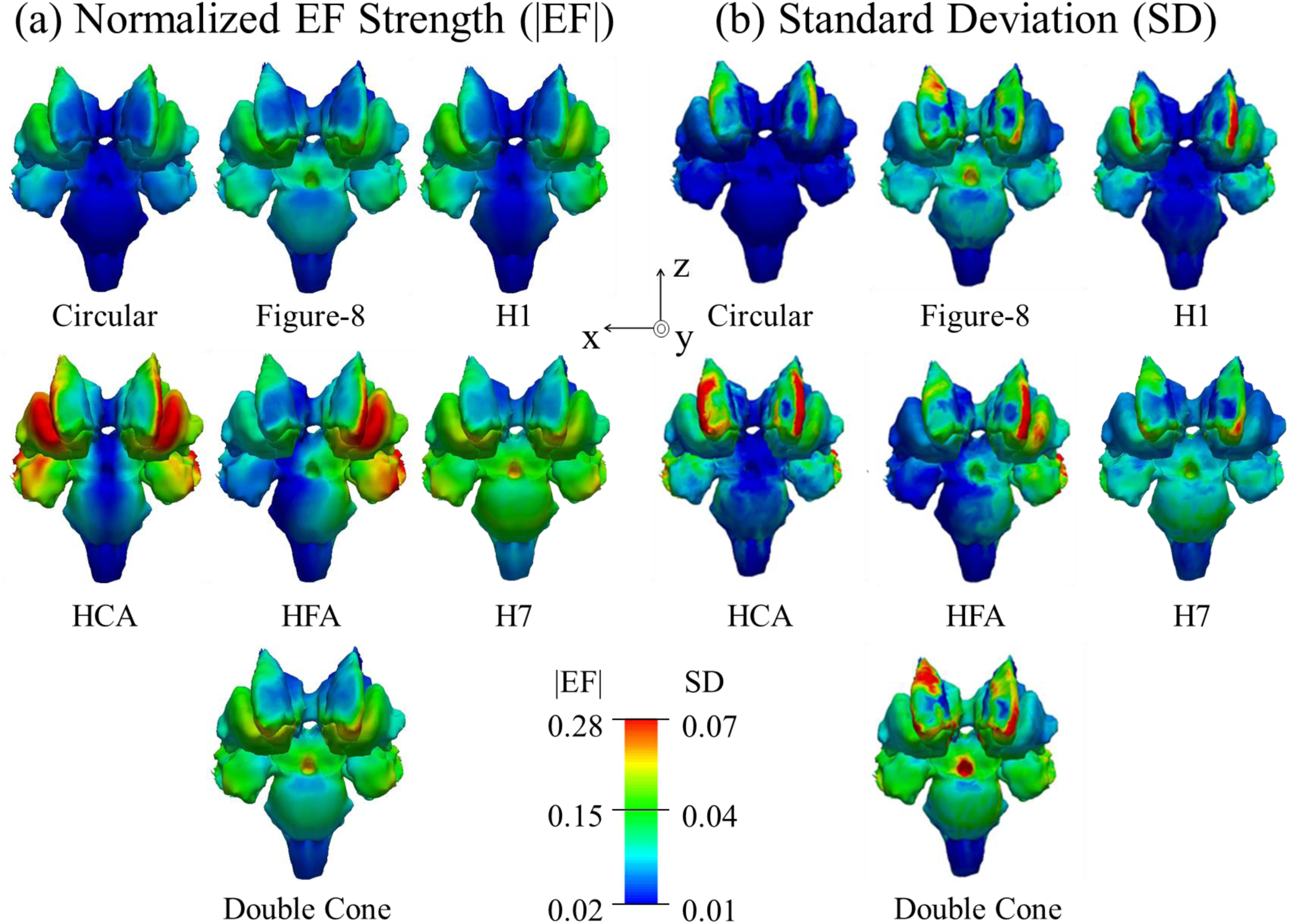
Group-level EF. (a) EF averaged over different models (*n* = 18) in the standard brain space of deep regions for the seven coils centered in cz position. (b) Relative standard deviation of the EF for each coil. Values are normalized by the maximum cortical EF.

### C. Trade-off between depth and spread and co-activation

Fig. 5(a) shows the tradeoff relationship between S_1/2_ and *d*_1/2_ considering field values normalized by maximum cortical EF. The ideal coil for dTMS would generate a high induced EF in deep regions (high *d*_1/2_) with small spread (small S_1/2_). The penetration depths using half-value metric correspond to the subcortical areas (< 2 cm from the cortical surface) and far from deep brain regions (> 4 cm in Table 2). HCA can reach high penetration and with a smaller spread. In general, double cone coil had a good trade-off between depth and spread; however far from deep brain regions.

**Figure 5.**
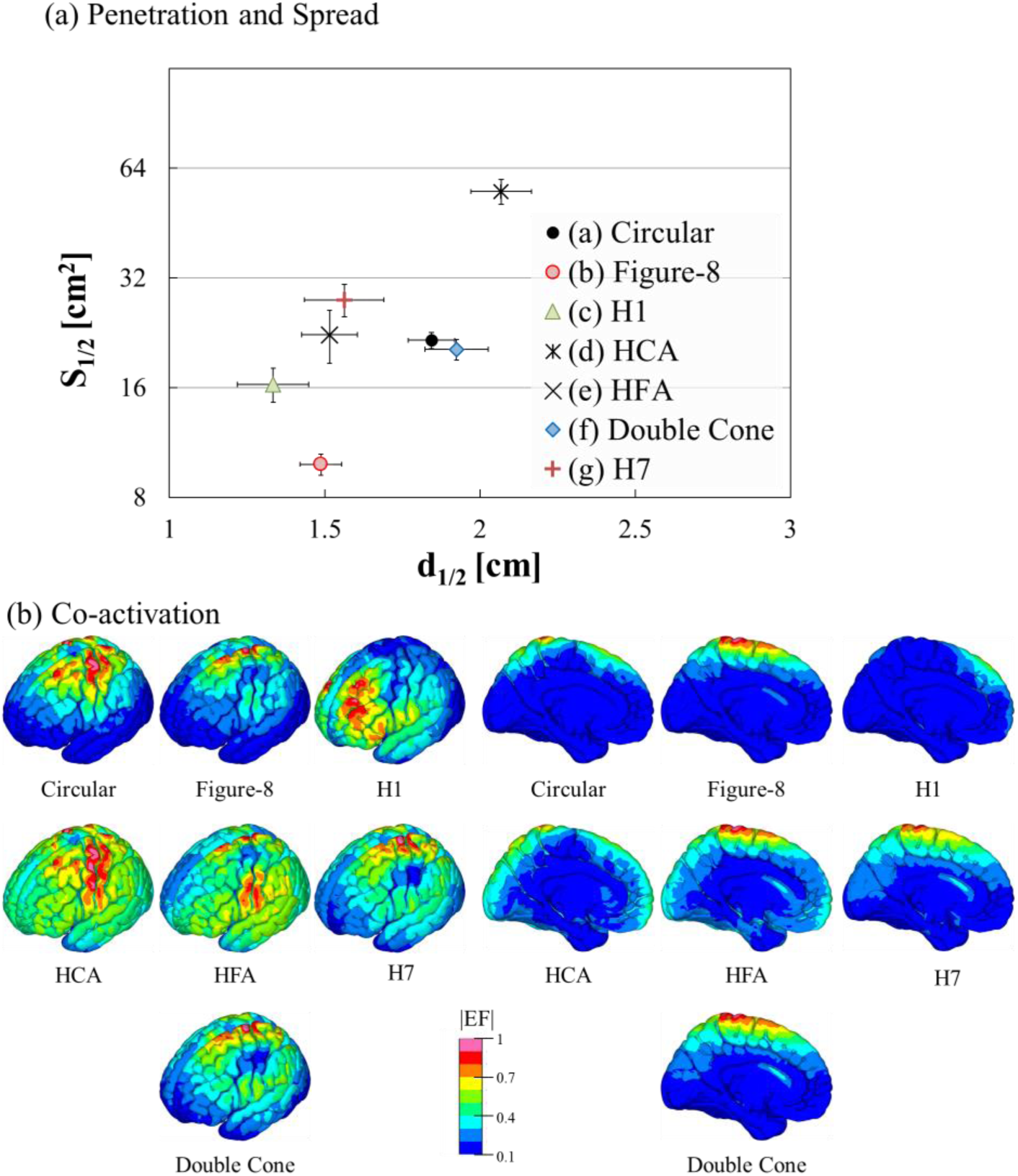
(A) Relationship between spread (S1/2) and depth from cortical surface (d1/2) of the EF (*n* = 18). (B) Group-level EF (*n* = 18) co-activation in the cerebral cortex and subcortical regions (EFs are normalized by the maximum value).

**TABLE 2.** Distance from the deep brain regions to the closest point of the scalp

| Deep brain structure | Distance<br>[cm] | Volume [cm<sup>3</sup>] |
| --- | --- | --- |
| Amygdala | $5.05 \pm 0.28$ | $2.33 \pm 0.06$ |
| Brainstem | $6.56 \pm 0.33$ | $4.39 \pm 0.14$ |
| Caudate | $4.69 \pm 0.22$ | $3.00 \pm 0.15$ |
| Hippocampus | $4.76 \pm 0.31$ | $3.15 \pm 0.16$ |
| NAcc | $5.47 \pm 0.30$ | $1.72 \pm 0.10$ |
| Putamen | $4.29 \pm 0.24$ | $3.56 \pm 0.20$ |
| Thalamus | $5.37 \pm 0.26$ | $3.94 \pm 0.16$ |

Co-activation of superficial brain regions is a consequence of the trade-off between depth and spread. A group-level map (*n* = 18) of the cortical and subcortical EFs is presented to quantify the effects of co-activation in Fig. 5(b). From the coils with higher EF percentage in deep brain regions (i.e., HCA and HFA,), co-activation of HCA is mainly in prefrontal and occipital areas, whereas HFA had large co-activation in cortical and subcortical regions below the cz position. Double cone had higher penetration than Figure-8 coil but also with considerable co-activation. Also, H1 is more suitable for focal activation of prefrontal cortex and underlying subcortical regions. H7 can activate deeper regions but with higher co-activation than H1.

### D. Optimized Group-level EF distribution

The coil position is optimized to maximize the group-level EF in different deep brain regions, as shown in Fig. 6(a). The coils are centered on scalp positions according to 10-10 EEG international system in all subjects. The results showed that the HCA has the highest group-level EFs (up to 50% in most of the deep brain regions), while double cone had the smallest group-level EFs (up to 30% in deep brain regions). Figure-8 also presents high EFs in hippocampus and amygdala relative to cortical EF; however, the dTMS-induced EF is limited by the TMS device power for figure-8 (Table A1, Appendix A). In contrast, double cone or H7 can reach deeper regions with higher EF at the expense of induced higher and wider spread electrical fields in superficial cortical regions. The optimal location of the Figure-8 coil is the temporal region, which is a closer location to deep regions than midline scalp positions, such as double cone and H7.

**TABLE A1.**
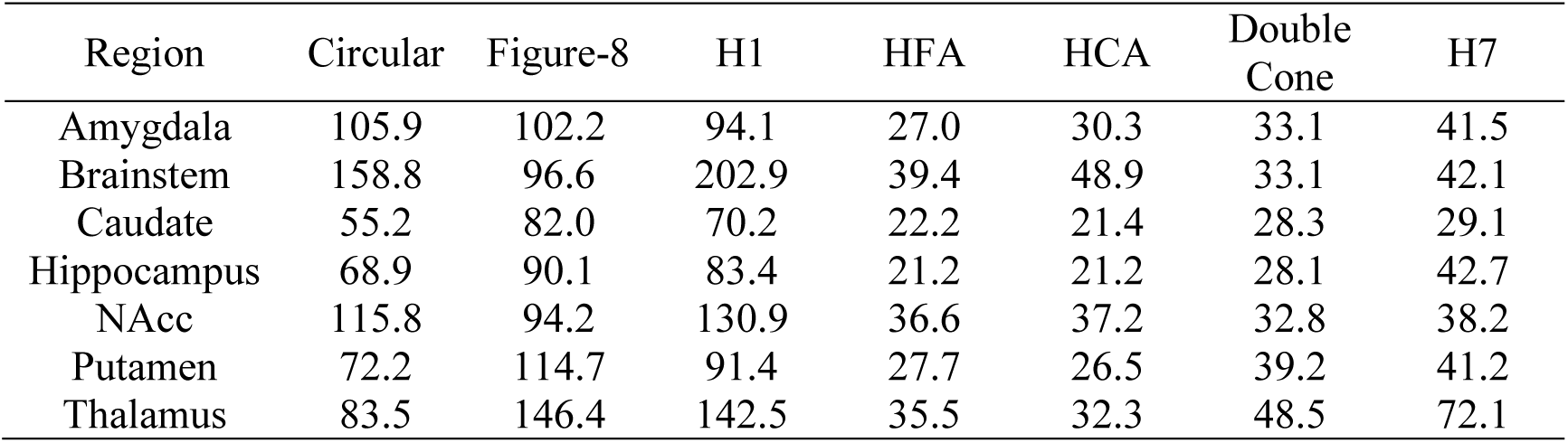
MSO Intensity (%) to generate 50 V/m at deep brain regions (*n* = 18)

**Figure 6.**
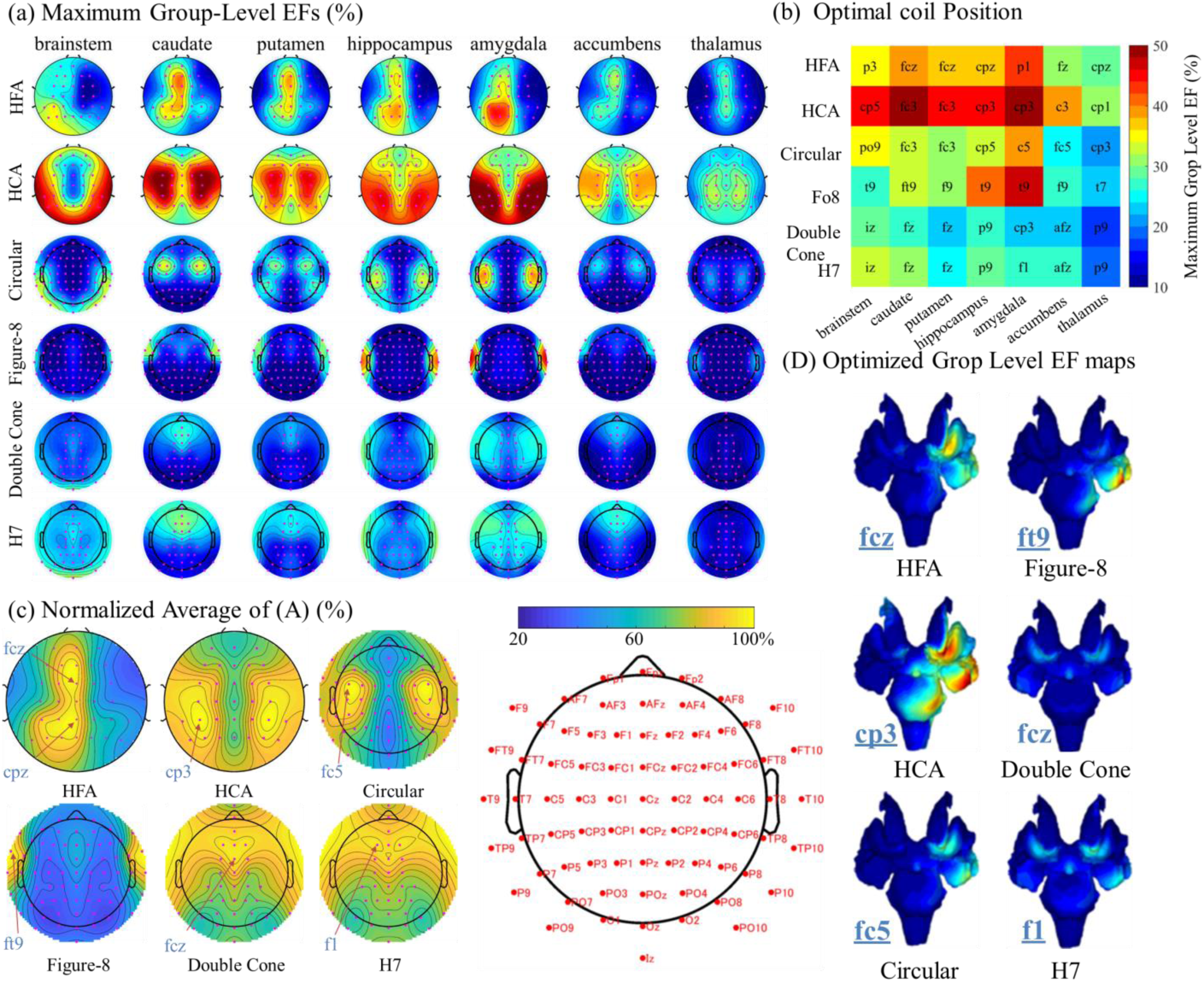
Optimized Group-level EF. (A) Scalp EF maps show the maximum group-level EF (*n* = 18) for individual deep brain region at different coils positions. (B) Coil position for maximum group-level EF at each deep brain region. (C) Average of normalized scalp maps for different depth regions. They represent optimized group-level EF to all deep brain regions. (D) Group-level maps optimized by individual deep regions (panel B) and for all deep brain region

In the case of the HCA, the optimal location is around fc3 for caudate, putamen, and NAcc, and around cp3 for the other deep regions. We also investigated the optimal coil location for all deep brain regions by averaging the normalized EF scalp maps across all deep regions for the same coil. We found that HFA (fcz or cpz), double cone (fcz), and H7 (f1) are optimal at medial scalp positions. In contrast, HCA (cp3), circular (fc5), and figure-8 (ftp) are optimal at more lateral scalp locations, as shown in Fig. 6(b).

## 4. DISCUSSION

We proposed a group-level EF approach in deep regions that allows generalizing dTMS coil performance in a group of subjects to overcome the limitation of using individualized head models to characterize coil performance in a population (Lu and Ueno 2017). For instance, Fig. 3(a) shows the variation of the EF distribution in deep regions of 18 subjects for the same coil according to subject anatomical differences in the head tissues. This agrees with previous studies where significant differences in the EF cortical distribution where found in the regions with higher individual neuroanatomical differences (Gomez-Tames et al. 2018). Reliable registration method was implemented to obtain group-level EF distributions on the standard space of deep brain regions to minimize subject differences (Fig. 3).

### A. dTMS coil characterization by Group-level EF

The dTMS coils were characterized by investigating the group-level EF distribution, trade-off between spread and depth, and co-activation. For fair characterization of the coil design, the coils were centered at the same location to reduce intra-subject differences (e.g., distance to deep regions if coils are placed at different locations in the same subject).

First, we observed consistent group-level EF hotspots on the deep brain region surfaces suggesting the possibility of targeting specific deep brain regions in Fig. 4(a). The HFA and HCA coils induced the highest EF in all deep brain regions. The highest EF was in the caudate for all coils. However, the other deep brain regions with maximum EFs distribution was different according to the coil. We also confirmed that the EF distribution of the HFA coil was asymmetric as the magnetic field was strengthened on one side, as this was the same direction of the current in the halo coil and one circular coil of the figure-8 coil. By contrast, the magnetic field on the other side was weakened due to the opposite effect.

Second, the most appropriate choice of coil settings should also be based on a well-balanced evaluation of penetration depth and focality. Fig. 5A confirmed the relationship between the penetration depth and spread was a trade-off (Deng et al. 2013; Gomez-Tames et al. 2018) based on half-EF value metric. The maximum penetration was smaller than 2 cm from the cortical surface and similar to (Deng et al. 2014); however, the group-average distance from the closest points between the scalp and each deep brain structure was approximately 4.3 to 6.6 cm, as shown in Table 2. Thus, the half-value metric is useful for evaluating EFs in subcortical regions but no suitable for EFs in deep brain regions. For the case of subcortical regions, double cone had a good trade-off between depth and spread.

Third, the concerns/implications of co-activation can be categorized into two groups: clinical side effects and mixed stimulation effects. Clinical side effects include transient headache or discomfort at the site of stimulation and seizure, the latter a major adverse event. A recent study (212 major depressive disorder (MDD) outpatients) showed that dTMS was safe with few and minor side effects apart from one seizure in a patient where a protocol violation occurred using H coil (Levkovitz et al. 2015). Mixed stimulation effects are generated due to co-activation of cortical and subcortical circuits with associated circuits to deep brain regions. Consequently, the conclusion derived from dTMS acting on a specific deep brain region should be taken with care and analysis of co-activation areas should also be considered, like the ones presented in this work (Fig. 5(b)). Therefore, the assumption that the stimulation of neighboring regions is acceptable needs to be further considered by the clinicians.

### B. Optimized dTMS localization by Group-level EF

To determine the maximum group-level EFs, scalp maps of EFs permitted to show the best locations in each deep region or whole deep regions for a population, as shown in Figs. 6(a) and 6(c), respectively. This method can be used to characterize and optimized dTMS coil in clinical applications where the same location is used in all subjects (one-for-all approach) usually based on 10-10 EEG international system. Optimized coil location induced a maximum group-level EF of 30-50% of the maximum EF value of the brain surface or 15-25% of the scalp surface. The coil with larger induced EF was the HCA and. Figure-8 also presented high EFs (45%) in hippocampus and amygdala for stimulation optimized for temporal location. One disadvantage of stimulation in the temporal region is that the pain perception threshold can be smaller than the parietal region. Even though optimized locations were explored, the EF levels (<50%) indicate that it is unlikely that dTMS can stimulate these deep areas without considerable side effects caused by the much stronger co-activation of other regions. In addition to the normalized EFs relative to the maximum on the cortex, we investigated the percentage of the maximum stimulation output (MSO) required to achieve similar EF levels in deep regions, as shown in Table A1 of Appendix A. The maximum MSO can be exceeded for some coils to target specific deep brain regions. HFA and HCA may require lower stimulation intensity to achieve similar EF in deep brain regions. Finally, this study used the medial-lateral coil orientation for optimization of the coil location as optimization based on coil orientation would have a negligible improvement in the dTMS-induced EFs for medial-lateral direction, as shown in Appendix B.

## 5. CONCLUSION

This study revealed the variability in EF distributions in deep brain regions resulting from inter-individual differences during dTMS. Despite these inter-individual variations, the proposed registration method for deep regions could determine a systematic tendency of the EF to derive the optimal coil for a group of subjects but with levels below 50% in comparison to cortical EFs for optimized coil localization. As a result, the first generalized map of targeted areas by different coils was presented for dTMS. These maps will enable the delivery of the most optimal dosage to the desired target, while also accounting for co-activation as an important factor to be included in dTMS studies. Future studies can use this approach to investigate new coil designs to facilitate maximum EF on specific deep structures and effects on different population segments (e.g., gender, age).

## APPENDIX A: MAXIMUM STIMULATION OUTPUT REQUIREMENTS

To induce the same level of EF in deep brain regions, each coil may require a different percentage of the maximum stimulation output (MSO) of the TMS device. Table A1 shows that the 100% MSO (corresponds to 160 A/μs) is exceeded for circular, Figure-8, and H coils when targeting specific deep brain regions for an induced EF strength of 50 V/m (Casali et al. 2010). The other coils are more suitable for achieving a larger field strength within the output capability of the TMS device.

## APPENDIX B: OPTIMIZED COIL ORIENTATION

The dTMS-induced EF variation due to coil orientation is investigated for HFA and figure-8 coil. Coil optimization for circular and HCA are not considered due to coil symmetry. Also, medial-lateral orientation is the optimal for double cone and H7 to fit the head shape. The coil is rotated from 0° (medial-lateral orientation) to 150° with steps of 30° in an anticlockwise direction (superior view of coil). The coil is located to the same scalp positions of Fig. 6A. The improvement of the EF on different regions is quantified by the percentage point variation taking as reference the EF for the medial-lateral orientation. Figure B1 shows no significant variation of the EF owing to coil orientation variation at the optimized coil position for deep brain regions (negligible for HFA and less than 6% percentage point for figure-8 at 120°).

**Figure B1.**
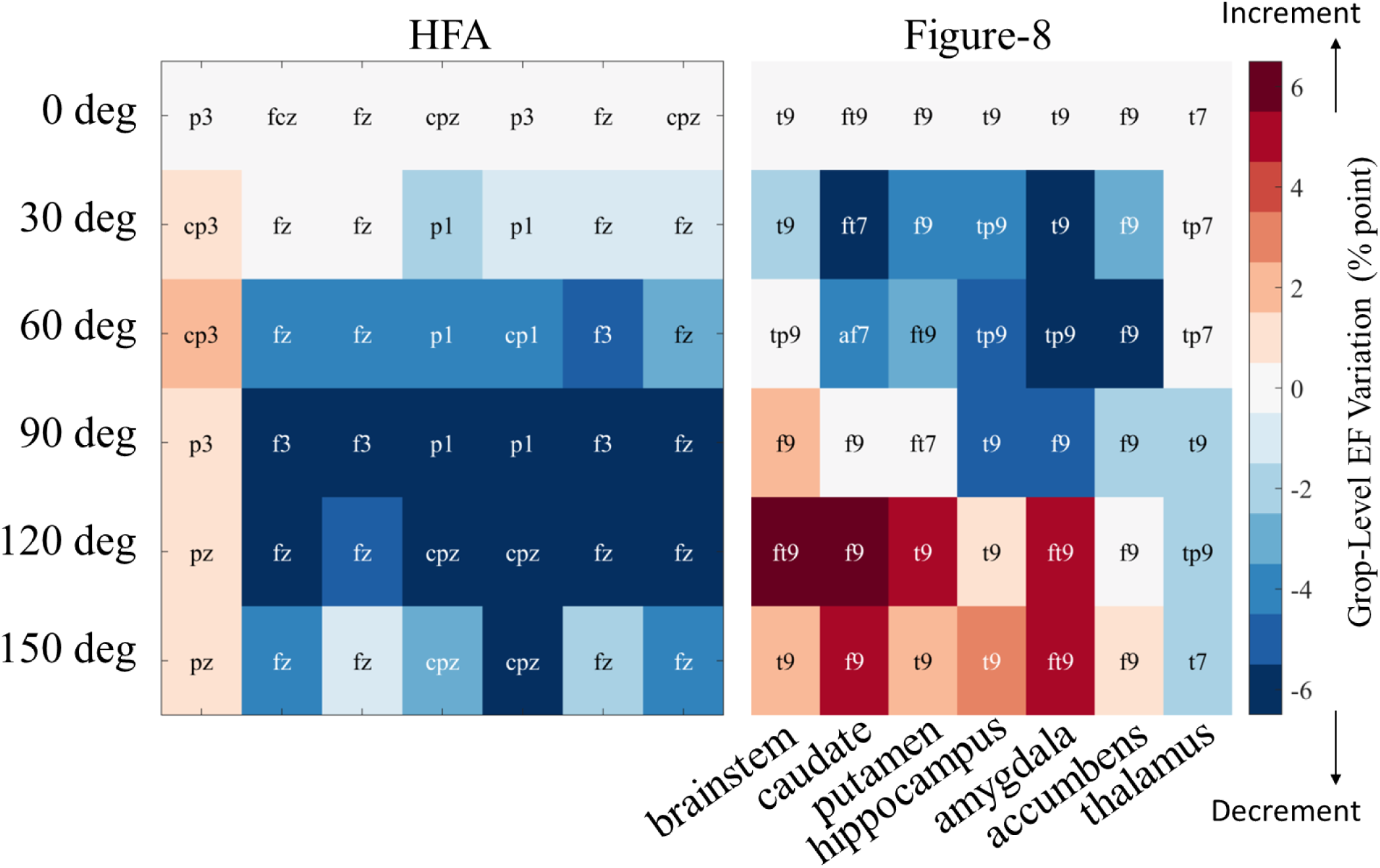
Coil angle variation effect. Group-level EF variation at optimized coil position considering angles from 0° to 150. The percentage point variation is respect to medial-lateral orientation (0°). Optimal coil positions are indicated (10-10 EEG system).

## ACKNOWLEDGMENT

This work was supported by a Grant-in-Aid for Scientific Research (A) (JSPS KAKENHI 17H05293) and a Grant-in-Aid for Early-Career Scientists (JSPS KAKENHI 19K20668) from the Japanese Society for the Promotion of Science (JSPS).

